# Variability in sampling of cortex-wide neural dynamics explains individual differences in functional connectivity and behavioral phenotype

**DOI:** 10.1101/2022.01.24.477572

**Authors:** Camden J. MacDowell, Brandy A. Briones, Michael J. Lenzi, Morgan L. Gustison, Timothy J. Buschman

**Affiliations:** Princeton Neuroscience Institute, Princeton University, Washington Rd, Princeton, NJ; Rutgers Robert Wood Johnson Medical School, 125 Paterson St, New Brunswick, NJ; Department of Psychology, Princeton University, Washington Rd, Princeton, NJ; Department of Anesthesiology and Pain Medicine at University of Washington, Seattle, WA; Department of Integrative Biology, University of Texas at Austin, Austin, TX

## Abstract

Individual differences in behavior are associated with changes in the correlation of neural activity between brain areas. Such differences in ‘functional connectivity’ are thought to reflect individual differences in brain structure that alter the flow of neural activity between regions. Here, in contrast, we show that individual differences in functional connectivity and behavior can be explained by differences in how frequently an individual expresses distinct cortex-wide spatiotemporal patterns of neural activity. This suggests variability in sampling of cortex-wide neural dynamics may underlie individuals’ unique behavioral phenotypes.

## Main

Behavior differs across individuals, ranging from typical to atypical phenotypes^1^. Understanding how individual differences in behavior relate to differences in neural activity is critical for developing individualized treatments of neuropsychiatric and neurodevelopmental disorders. Considerable work has linked behavioral variability to differences in how correlated neural activity is between brain areas, termed ‘functional connectivity.’ For example, patterns in functional connectivity vary across individuals^2^ and are disrupted in autism^3^, schizophrenia^4^, and depression^5^. However, the changes in neural activity that underlie this altered functional connectivity remain unclear.

The prevailing hypothesis is that individual differences in functional connectivity, and, in turn, behavior, reflect changes in the ability of neural activity to flow between brain areas^6–10^. This has been hypothesized to reflect changes in the anatomical connectivity between brain regions^11^ or changes in the ability of brain regions to synchronize^12^. Here, we present evidence for an alternative hypothesis: differences in which patterns of neural activity are expressed (i.e., ‘sampled’) contribute to individual differences in functional connectivity. This argues that differences in functional connectivity are not due to changes in anatomy restricting how neural activity can flow through the brain, but rather changes in the mechanisms that select which pattern of neural activity are expressed at each moment in time.

To understand how individual differences in behavior relate to changes in cortex-wide neural dynamics, we combined behavioral assays and mesoscale imaging in mice^13^. Since we were interested in studying a heterogenous population of mice with a spectrum of individual differences, recordings were done in ‘typical’ (wildtype) mice and in mice exposed to valproic acid (VPA) *in utero* (Fig. 1A; see Methods). VPA mice exhibit altered social and repetitive motor behaviors^14^, and are widely used to study how changes in neural structure and function relate to behavior^15–18^ (often as a model of cognitive dysfunction, including autism^19,20^). Importantly, VPA exposure provides an etiologically valid way of inducing a spectrum of behaviors in an otherwise genetically identical cohort of mice (n=52 mice; 27 VPA, 25 saline controls).

**Figure 1:**
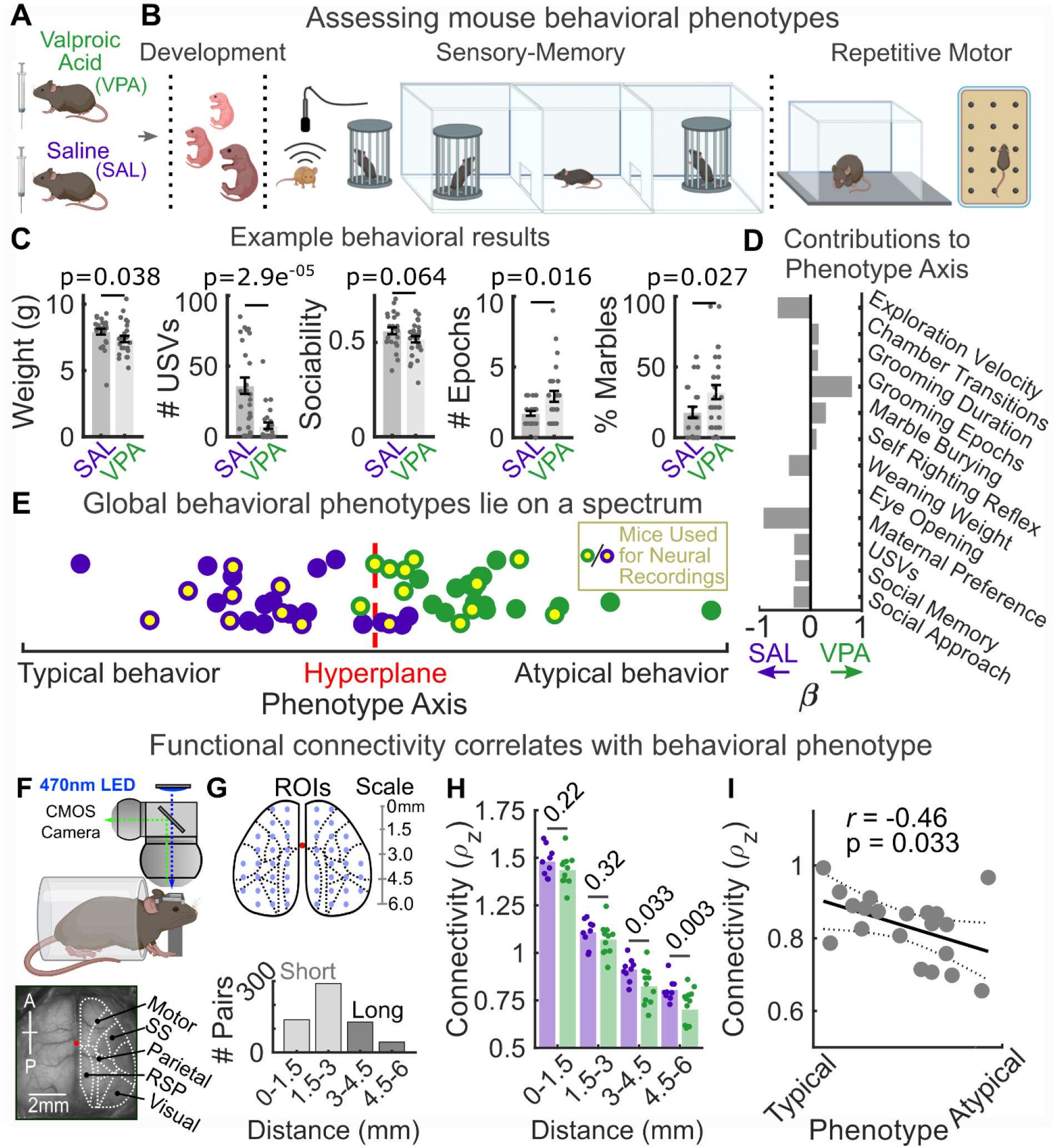
Cortical functional connectivity correlates with behavioral phenotype in mice. **(a)** Schematic of the *in utero* valproic acid (VPA) model used to investigate a diversity of behavioral phenotypes **(b)** Schematic of example behavioral assays. From left to right: weaning weight and developmental milestones, pup ultrasonic vocalizations (USVs) in response to maternal separation, three-chamber social approach and social novelty, self-grooming behavior, and digging behavior in a marble burying assay. See Methods for complete list of assays and measures. **(c)** Example results comparing behavior of VPA and saline-exposed (SAL) control animals. From left to right: animal weight at weaning, number of pup USVs in response to maternal separation, animals sociability score measured from social novelty assay, number of self-grooming epochs, percent of marbles buried in marble burying assay. See methods for details and Extended Data Figure 1 for complete results. **(d)** Beta weights of behavioral measures

Animal behavior was assessed with a battery of 9 tests targeting 12 measures of general developmental, sensory-memory, and motor behavioral domains (Fig. 1B; see Methods). Consistent with previous literature^14,19^, individual tests showed significant differences between the saline-exposed controls and VPA mice (Fig. 1C and Extended Data Fig. 1A-C). Consistent with a spectrum of individual behavioral phenotypes^1^, there was considerable variability across individuals, both between and within groups. To quantify this spectrum, we projected the multivariate behavioral data onto a single ‘phenotype axis’. The phenotype axis was defined as orthogonal to the linear hyperplane that maximally separated the behavior of VPA and saline mice (across all measures; Fig. 1D-E; see Methods). Projecting each animal’s behavior onto this axis captured where that individual fell along the behavioral spectrum. For example, some VPA mice demonstrated more typical behaviors than others, while some saline animals showed more atypical behavioral features. Similar degrees of behavioral diversity were observed in each behavioral domain (Extended Data Fig. 2A-B).

To study cortex-wide neural dynamics, we used mesoscale calcium imaging to record activity of excitatory neurons across the dorsal cortex of a subset of VPA (n=11) and saline (n=9) mice (Fig. 1F; all Thy1-GCaMP6f^21^ mice, see Methods). These animals spanned the behavioral spectrum of the larger population (Fig. 1E, yellow dots). Motivated by previous work^18,22^, we tested whether functional connectivity varied across animals. Functional connectivity was measured as the zero-lag pairwise correlation in neural activity between a grid of sites across the dorsal cortex (Fig. 1G). We found that functional connectivity between local cortical regions (<3mm) was similar between groups, but long-range (>3mm) connectivity was significantly decreased in VPA animals relative to saline controls (Fig. 1H). Reflecting its relationship with behavior, the difference in long-range connectivity was significantly negatively correlated with phenotype across animals (Fig. 1I; *r*=-0.46 p=0.033 permutation test, n=20 mice). These results are consistent with the spectrum of behavior and functional connectivity seen in humans^2–5^, highlighting the face validity of the VPA model.

As detailed above, the predominant hypothesis is that these differences in functional connectivity reflect changes in how neural activity flows across the cortex. To test this, we quantified the spatiotemporal dynamics of cortical activity by decomposing cortex-wide neural activity into a set of 16 ‘motifs’ (schematized in Fig. 2A; see Methods and Extended Data Fig. 3A-G). Each motif captured a unique cortex-wide spatiotemporal pattern of neural activity that lasted for ~1 second and is thought to reflect a specific cognitive process^23^. For example, Motif 12 is a cortex-wide wave of neural activity originating in somatosensory and motor areas and travelling posteriorly to retrosplenial and visual cortices; which is thought to coordinate neural activity across areas^24,25^. In contrast, Motif 5 is a spatially static, bilateral, burst of activity in the visual cortex, associated with the processing of visual stimuli^23,26^. By tiling the 16 unique motifs over time, we could accurately reconstruct cortical activity in all mice (Fig. 2B; see Methods, and Extended Data Fig. 4 and Extended Data Table 1 for all 16 motifs).

**Figure 2:**
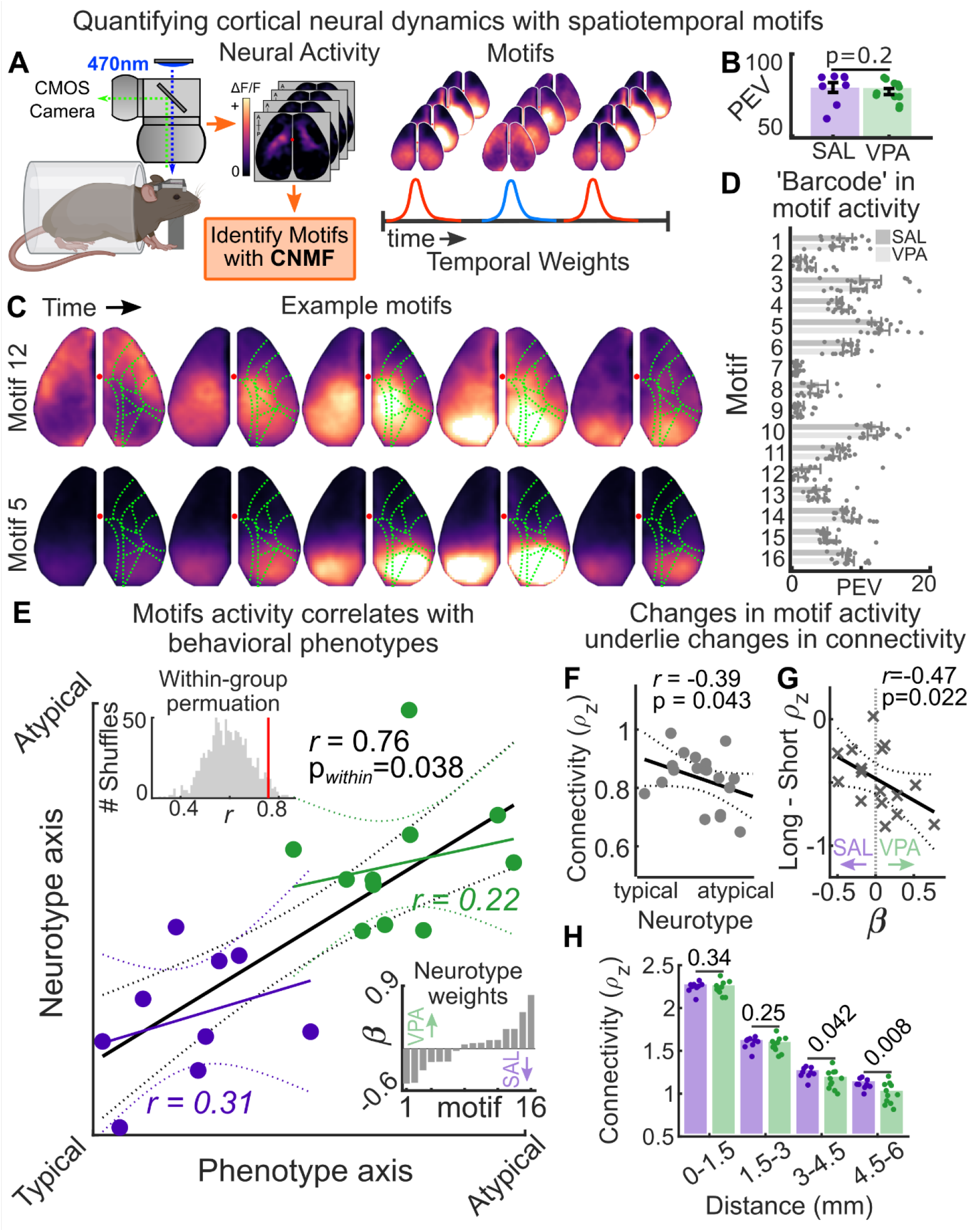
Spatiotemporal motifs in neural activity correlate with behavioral phenotype and explain the relationship between functional connectivity and behavior. **(a)** Schematic outlining quantification of spatiotemporal motifs in cortical activity. (left) Neural activity was decomposed with convolutional matrix factorization into (right, top) recurring motifs and (right, bottom) their temporal weightings. **(b)** Percent variance in cortical neural activity explained by 16 spatiotemporal motifs shared across all animals. Mean saline: 79.59% CI: 70.88%-83.80%; VPA: 77.12%, CI:72.80%-80.38%; no difference between groups, p=0.20; Mann-Whitney U-Test and no correlation with behavioral phenotype; *r*=0.031, p=0.54, permutation test. **(c)** Example motifs. Green lines outline anatomical parcels as in Figure 1E. Red dot denotes bregma. **(d)** ‘Barcode’ in the relative percent variance in neural activity explained by each motif per mouse. **(e)** Correlation between neurotype axis and phenotype axis. Top inset shows correlation values from random within-group permutations. Red line shows observed correlation. Bottom inset shows beta weights of motif contributions to the neurotype axis. **(f)** Strength of animals’ long-range functional connectivity compared to neurotype. **(g)** Within-motif long-range minus short-range connectivity compared to neurotype axis beta weights (from Fig. 2E). **(h)** Strength of functional connectivity as a function of distance between groups in data reconstructed from motif activity. Long range functional connectivity differed between groups: for 3-4.5mm *ρ*_z_ saline=1.24 CI: 1.19-1.26; VPA= 1.16 CI: 1.11-1.23 p=0.042; for 4.5-6mm saline=1.11 CI: 1.07-1.15; VPA=1.00 CI: 0.94-1.08 p=0.008; permutation test. Data points reflect individual mice in panel B,D,E,F, and H and motifs in panel G. All errorbars show mean +/- SEM. Solid and dotted lines in panels E,F,G show least squares fit and 95% confidence bounds, respectively.

Given that motifs reflect spatiotemporal patterns in neural activity across brain areas, we can use them to quantify whether the dynamics in neural activity changes between individuals. If changes in functional connectivity are due to altered patterns in the flow of neural activity across the cortex, then the spatiotemporal patterns captured by motifs will be different between individuals. In contrast, if changes in functional connectivity are due to variability in how motifs are sampled, then all mice should share the same motifs, but how often a motif occurs will differ across animals. Consistent with a change in sampling, the same 16 motifs captured over 75% of the variance in cortical neural activity in both VPA and saline animals (Fig. 2B; no difference between groups). In addition, motifs independently identified in each group of animals were also the same: linear decoders were unable to differentiate between motifs from VPA or saline animals (mean decoding accuracy across motifs=51.46%, CI 48.37%-54.81%, p=0.34, Wilcoxon Signed-Rank Test; see Methods). This was not due to an inability of linear decoders to discriminate motifs; they easily discriminated between the 16 motifs (accuracy=99.08%, CI 98.98%-99.18%, p<0.0001, Wilcoxon Signed-Rank Test).

The similarity of motifs across groups suggests disruptions in functional connectivity are not due to altered patterns of cortical neural activity. Instead, what differed between groups was how frequently a motif was expressed. In each mouse, different motifs were active more/less often, and so contributed to more/less of the variance in neural activity (see Methods). This motif activity was heterogenous across animals, and no single motif clearly separated VPA and saline groups (Fig. 2D; no differences significant at p=0.05, Mann-Whitney U-Test, although, in general, the likelihood of expressing a motif was more uniform in VPA mice; see Extended Data Fig. 5). However, by measuring the relative variance captured by each motif, we could create a ‘barcode’ of neural dynamics for each animal. Using these barcodes, a cross-validated linear classifier could accurately classify VPA and saline animals (accuracy=86.61%, CI 80.00%-95.00%; see Methods). The beta weights of the classifier were distributed across all 16 motifs, supporting the conclusion that discriminability between groups relied on global changes in sampling, rather than disruption of any single motif (Fig. 2E). Similar to behavior, we could estimate each animal’s ‘neurotype’ by projecting its individual barcode onto the axis orthogonal to the hyperplane that best separated the two groups (Fig. 2E; see Methods). Despite being defined independently and measured weeks apart, individuals’ neurotype and phenotype were significantly positively correlated (Fig. 2E; *r*=0.76, *p*=0.038, within-group permutation test). In other words, animals with more atypical expression of motifs also had more atypical behavior. Deeper analyses confirmed that animals’ neurotype was broadly associated with individual behavioral tests; but as expected given the dominance of motor representations in cortical activity^26,27^, neurotype was most strongly related to tests of motor phenotype (Extended Data Fig. 6). Importantly, control analyses ruled out potential confounds, such as differences in motor activity during recordings, brain size, GCaMP6f expression, neural variance, and confirmed that motif activity was stable across days and that Thy1-GCaMP6f mice did not exhibit epileptiform activity^28^ (Extended Data Fig. 7).

Differences in the distribution of motif activity explained differences in functional connectivity across animals: an animals’ neurotype was significantly correlated with long-range functional connectivity (Fig. 2F; *r*=-0.39 p=0.043, permutation test). Furthermore, removing neurotype contributions to functional connectivity eliminated the correlation between connectivity and phenotype, supporting the hypothesis that motif activity drove this relationship (*r*=-0.17 p=0.24, permutation test). In addition, the strength of long-range functional connectivity relative to short-range functional connectivity within a motif was negatively correlated with the motifs’ neurotype beta weights (Fig. 2G; *r*=-0.47 p=0.022, permutation test, n=16 motifs). In other words, motifs capturing patterns of neural activity with stronger long-range connectivity were more active in ‘typical’ animals and motifs with weaker connectivity were more active in ‘atypical’ animals, explaining the observed changes in functional connectivity. Finally, to test whether changes in motif activity were sufficient to explain the observed changes in functional connectivity, we reconstructed cortical activity of all animals using the shared set of 16 motifs. These reconstructed neural dynamics showed a similar decrease in long-range functional connectivity as the original data (Fig. 2H). By using the same motifs in all animals, these changes in functional connectivity can only arise from changes in the expression of motifs, and not differences in the spatiotemporal structure of motifs across animals.

Our results suggest that individual differences in behavior (and functional connectivity) are due to individual variability in the expression of motifs. Given that motifs capture patterns of neural activity associated with specific cortical processes^23^, one hypothesis is that a set of control mechanisms may ‘sample’ motifs to engage in context-appropriate cognitive processes. If true, then motif activity should adapt to different behavioral contexts. To test this, we compared motif activity between two contexts: when mice were alone (as above) and when mice were interacting with a second animal in a stimulus-enriched environment (‘paired’ context; Fig. 3A; see Methods). Consistent with the hypothesis that motifs are adaptively sampled, the expression of motifs was significantly different between contexts in all mice (all p-values <0.001; see Methods).

**Figure 3:**
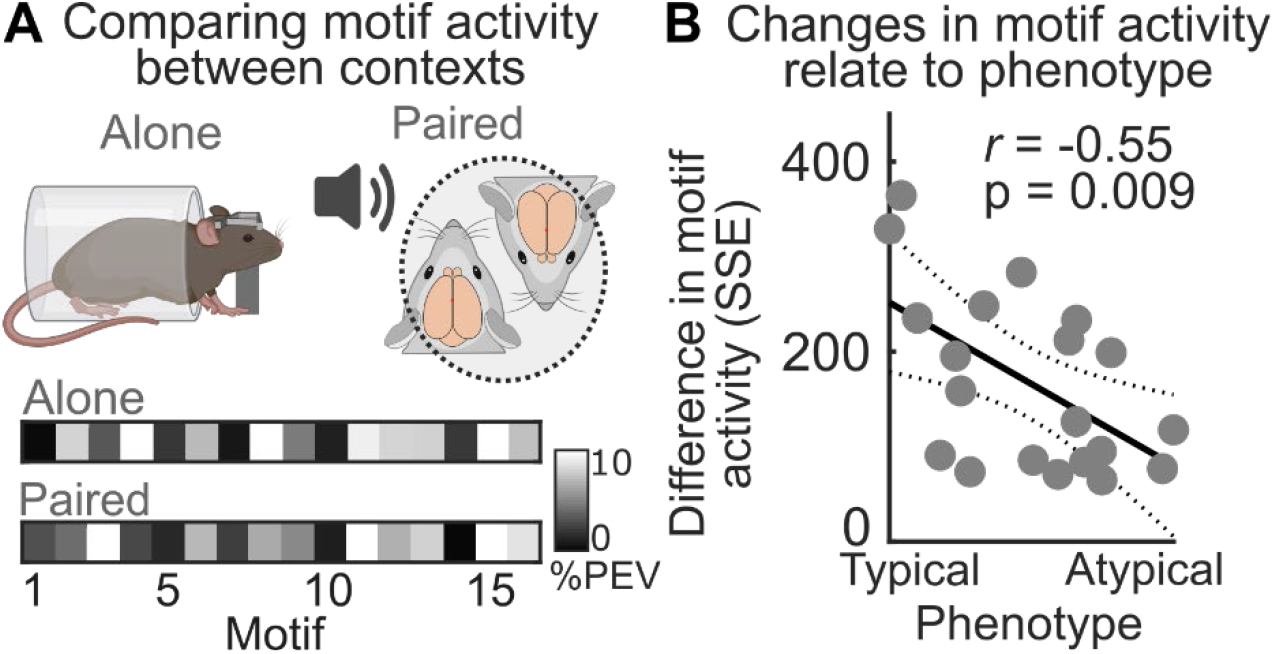
Individual variability in motif sampling across behavioral contexts relates to behavioral phenotype. **(a)** Top: Schematic of contexts. Bottom: In each context, motif activity was measured as the barcodes in the percent explained variance (PEV) of cortical neural activity captured by each motif. **(b)** Magnitude of change in motif activity between context compared to behavioral phenotype. Change in motif activity was measured as the sum squared error between motif barcodes in each context. Data points show individual mice. Solid and dotted lines show fit, and 95% confidence bounds, respectively.

However, the magnitude of change in motif activity between contexts was significantly negatively correlated with behavioral phenotype across mice (Fig. 3D; *r*=-0.55 p=0.009, permutation test). In other words, more atypical mice showed less of a change in motif activity to the novel, stimulus-rich paired context, suggesting an inability to adapt motif expression to match new contexts. Consistent with this, how much an individual’s motif activity changed between contexts was most strongly related to their performance on behavioral tests involving novel contexts and/or social stimuli (Extended Data Fig. 8). Furthermore, the sequence of motifs over time was more regular in behaviorally atypical mice, suggesting they were more rigid in the ordering of motif expression (Extended Data Fig. 9). Together, these results support the hypothesis that individual differences in behavior are, at least partially, due to variability in the control or selection of motif expression.

In sum, our results suggest that individual differences in behavior and functional connectivity are driven by changes in how frequently specific spatiotemporal motifs in cortical activity occur, rather than the gain or loss of specific motifs or changes in the spatiotemporal patterns of neural activity within motifs. In particular, behavioral phenotype was correlated with how well an individual adapted motif expression to a change in the environment, suggesting that individual differences may reflect differences in the mechanisms that control the sampling of motifs. This likely depends on a distributed set of mechanisms, including top-down cognitive control, neuromodulators, and internal state (e.g., homeostatic mechanisms). Our results set the stage for future work investigating how biases in sampling of neural population dynamics may manifest to drive cognitive changes in health and disease.

## Acknowledgments

We thank the Buschman lab for their detailed feedback during the writing of this manuscript and the Princeton Laboratory Animal Resources staff for their support. This work was funded by a grant from SFARI 670183 (T.J.B.), NIH DP2 EY025446 (T.J.B), NIH NCATS Award TL1TR003019 (C.J.M), and T32 MH065214 (M.L.G).

## Author Contributions

Conceptualization: TJB and CJM; Experimental Design: TJB, CJM, MLG; Surgeries and Data Collection: CJM, BAB, MJL; Analysis and Visualization: CJM; Writing: CJM and TJB; Editing; TJB, CJM, BAB, MJL, MLG Funding Acquisition: TJB, C.J.M, M.L.G; Supervision: TJB

## Material and Methods

### Data and Code Availability

Example data and figure generation code will be available upon acceptance on our GitHub repository (https://github.com/buschman-lab). Raw data is available upon reasonable request.

### Animal Care and Details

All experiments and procedures were carried out in accordance with the standards of the Animal Care and Use Committee (IACUC) of Princeton University and the National Institutes of Health. Behavioral testing took place post-natal days (PND) 5-45, surgery and habituation PND 45-55, solo imaging PND 60-70, and paired imaging PND 80-100. Mice were group housed with same sex littermates prior to surgery and single housed post-surgery on a reverse 12-hr light cycle. All experiments were performed during the dark period, typically between 12:00 and 18:00. Animals received standard rodent diets and water *ad libitum.* Both male (n=23) and female (n=29) mice were used. All mice were offspring of 8 liters resulting from crosses between C57BL/6J and C57BL/6J-Tg(Thy1-GCaMP6f)GP5.3Dkim/J mice. 5 litters were exposed to valproic acid (VPA) *in utero* and 3 with saline controls. Offspring (VPA=27; saline=25) were used for behavioral experiments. Thy1-GCaMP6f heterozygous animals (VPA=11; saline=9) were also used for neural recordings. The first day of imaging in each animal was screened for potential epileptiform events associated with GCaMP6f lines, and none were observed, consistent with previous work expressing GCaMP under the Thy1 promoter^28^ (Extended Data Fig. 7A). Histology on 8 VPA and 5 saline animals confirmed similar levels of GCaMP6f expression between groups (Extended Data Fig. 7B). Data from the 9 saline animals used for imaging has been reported in a previous study^23^.

### *In Utero* Valproic Acid Model

The *in utero* VPA mouse model followed the procedures used in previous work^29^. Pregnant dams were administered a single 600mg/kg subcutaneous injection of valproic acid sodium salt (Sigma Aldrich) dissolved in sterile water on embryonic day 12.5. Saline (control) litters received equivalent volume of saline.

### Behavior and Development Testing

A battery of 9 behavioral tests was used to quantify animal behavior and general development. Tests, ordered by their post-natal day (PND), are reported below. The experimenter was blind to animal groups throughout behavioral testing and quantification of the behaviors. Experimental setups were cleaned with ethanol and allowed to dry between animals to remove olfactory cues. See Behavioral Measures subheading below for quantification details.

#### Maternal Separation Ultrasonic Vocalizations

PND 5. Pups were assessed for isolation-induced ultrasonic vocalizations (USVs)^30^. Tests were performed in a dark, quiet testing room after the litter had been habituated to the room for 10 minutes. The cage was kept on a warming pad throughout testing. Pups were removed from the litter and placed in a plastic dish within a sound-attenuating chamber 5cm below an ultrasonic detector (Dodotronic Ultramic200K, Dodotronic; Part # UM200K). USVs were recorded for 3 minutes. Data was acquired using Audacity software on a PC (Windows 10). USV number was quantified using MUPET^31^. Pup axillary temp was recorded prior to being replaced in the home cage. No difference in temperature was noted between groups (difference between means of each group = 0.57°F, p=0.89; Mann-Whitney U-Test).

#### Self-righting reflex

PND 5-14. Pups were placed on their backs and given 10 seconds to self-right. This was repeated three times. Reflex was considered developed if the pup righted all times.

#### Eye Opening

PND 12-18. The number of eyes open was recorded. Partially open eyes were considered open.

#### Three Chamber Maternal Scent Preference

PND 14. A 11” x 8” plastic box was split into thirds. The lateral thirds were filled ~5mm deep with bedding from the pup’s cage (familiar) or the cage of a stranger litter. A pup was placed in the center third and the time spent in either side third was manually recorded for a duration of 60 seconds. An animal was considered in a given third if its nose was in that region. This was repeated 3 times. The cage was rotated 180 degrees between each repeat to control for side preference.

#### Three Chamber Social Approach and Social Novelty

PND 40-45. Experiments were performed in a dimly lit room. Data was recorded using a video camera (Logitech Brio). *Habituation*: The test animal was habituated to a 23” x 14.75” custom three chamber polycarbonate box for 5 minutes (lateral chambers = 7.75” x 14.74”, Middle chamber 9” x 7.5”). *Social Approach*: After 5 minutes, the animal was secured in the center chamber and a metal cage was introduced to each lateral chamber. One cage contained a stranger animal, the other was empty. The test animal was released into the middle chamber, and activity recorded for 10 minutes. *Social Novelty*: After 10 minutes the test animal was secured in the center chamber and a new stranger mouse was added to the previously empty cage. The test animal was released, and activity recorded for 10 minutes. Stranger animals were age and sex matched and were randomly sampled from a pool of ~4-8 animals for each test mouse (with replacement). Sides were rotated between test mice. Animal activity was automatically quantified using custom MATLAB scripts that tracked the animals’ centroid (e.g., abdomen) over time.

#### Weaning Weight

PND 20. Animals were weighed.

#### Marble Burying Assay

PND 40-45. Performed in a dimly lit room. The test animal was habituated to a clean home cage with 1.5-inch-deep fresh bedding for 15 minutes. The animal was then removed from the cage and 14 marbles were placed on top of the leveled bedding in a grid like pattern. Animals were returned to the cage for 15 minutes after which the number of marbles >75% covered by bedding was recorded.

#### Repetitive Grooming Assay

PND 40-45. Experiments were performed in a dimly lit room. The test animal was habituated to a clean home cage with no bedding for 15 minutes. Grooming behavior was then manually scored (duration and number of separate epochs) for 10 minutes.

### Surgical Procedures

Surgical procedures followed previous work^23^. Mice were anesthetized (isoflurane: 1.5%) and administered analgesics (Buprenorphine, 0.1mg/kg; Meloxicam, 1mg/kg) and provided subcutaneous fluids (sterile saline, 0.01mL/g). The dorsal scalp was shaved and disinfected with betadine and 70% isopropanol. The dorsal cranium was exposed, and the periosteum removed. The dorsal cranium was rendered optically accessible by applying a thin layer of clear dental acrylic (C&B Metabond Quick Cement System) polished with a rubber rotary tool tip (Shofu, part #0321; Dremel, Series 7700) and coated with clear nail polish (Electron Microscopy Sciences, part #72180). A custom titanium headplate with a 11mm trapezoidal window was cemented to the skull to permit head fixation. Mice recovered in a clean home cage and received post-op analgesia (Meloxicam, 1mg/kg 24 hours post-surgery).

### Widefield Imaging

Widefield imaging followed previous work^23^. Mice were head-fixed in a 1.5 x 4-inch polycarbonate tube under a custom-built fluorescence macroscope consisting of back-to-back 50 mm objective lens (Leica, 0.63x and 1x magnification), separated by a 495nm dichroic mirror (Semrock Inc, FF495-Di03-50×70). Excitation light (470nm, 0.4mW/mm^2^) was delivered through the objective lens from an LED (Luxeon, 470nm Rebel LED, part #SP-03-B4) with a 470/22 clean-up bandpass filter (Semrock, FF01-470/22-25). Fluorescence was captured in 75ms exposures (FPS = 13.3Hz) by an Optimos CMOS Camera (Photometrics). The macroscope was focused below superficial blood vessels, approximately 500um below the skull surface.

Mice were habituated to head fixation for ~5 minutes and then imaged for 12 minutes. Images were acquired as three, 4-minutes stacks of TIFF images at 980×540 resolution (~34um/pixel) using Micro-Manager software (version 1.4) on a PC. Mice were imaged for 4-6 consecutive days for a total of 105 recordings.

For paired imaging, animals were positioned in individual polycarbonate tubes. The animals faced one another, approximately eye to eye along the anterior-posterior axis and equidistant from the macroscope objective along the z axis. Animal snouts were separated by 5-7mm; close enough to permit whisking and social contact but prevent adverse physical interactions. A 1mm plexiglass divider at snout level ensured no paw/limb contact. To permit a wider (~30×20mm) field of view, the macroscope objectives were replaced with 0.63x and 1.6x magnification back-to-back objectives (lens order: mouse, 0.63x, 1.6x, CMOS camera). Images were acquired at 1960 x 1080 resolution (~34um/pixel) for 12 consecutive minutes. Each pair of animals was recorded for 3 consecutive days.

Mice pairs were age-matched, primarily non-littermate, and included same sex, opposite sex, VPA-VPA, VPA-saline, and saline-saline pairs. Each animal participated in 1-3 pairs over the course of 2-3 weeks with at least 2 days between new pairings, for a total of 165 sessions across all animals. Manual inspection identified 28 sessions in which whiskers of one animal entered the imaging field of view of the paired animal; these sessions were excluded from further analysis, yielding 137 sessions in total. Throughout imaging, animal pairs were provided with auditory stimuli consisting of naturalistic adult ultrasonic vocalizations, synthetic vocalizations, or ‘background’ noise; the details of which are supplied in our previous manuscript^23^.

### Histology

Mice (n=13; saline=5, VPA=8) were transcardially perfused with 4.0% paraformaldehyde (PFA), brains were dissected and postfixed. Histochemistry was carried out on 50μm-thick free floating coronal brain sections and incubated in primary antibodies against green fluorescent protein (GFP) and counterstained with DAPI (Extended Data Fig. 7B). Sections were imaged with a Zeiss confocal microscope LSM 700 (solid-state lasers: UV 405; argon 458/488; Oberkochen, Germany). For cell density analysis sections were equally sampled to allow for cross comparisons and imaged at 20x, taking z-stacks at 2.0μm intervals. All slides were coded prior to analysis. All images were analyzed using FIJI.

### Statistical Analysis

All analyses were performed in MATLAB (Mathworks). No statistical methods were used to pre-determine sample sizes. The number of mice used in the study was based on previously published widefield imaging studies^32,33^. All statistical tests, significance values, n values, and associated statistics are denoted in the main text or figure legends. Unless otherwise noted, hypothesis tests were one-tailed. Permutation tests were run using 1000 permutations. 95% confidence intervals were computed with 1000 bootstrapped samples. Correlation values were Fisher r-to-z transformed prior to computing secondary statistics (i.e., mean and confidence intervals). Unless otherwise noted, behavioral measures were taken from distinct samples (i.e., mice) while neural measures, such as motifs, were taken as the average of repeat measures in the same mouse (i.e., per recording epoch).

### Behavioral Measures

Behavioral measures were quantified as follows:

#### Sensory-Memory

##### Maternal Separation Ultrasonic Vocalizations

Measured as the total number of USVs emitted.

##### Maternal Scent Preference

Measured as the 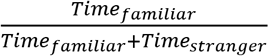 across the three runs.

##### Social Approach Index

Same as maternal scent preference but comparing the time in the compartment of the stranger mouse versus the compartment with the empty cage. Social approach index was bias-corrected for side preference by subtracting the magnitude of preference of the stranger animal compartment during habituation (relative to no bias: i.e., habituation bias - 0.5)

##### Social Novelty Index

Same as maternal scent preference but comparing the time in the compartment of the ‘familiar mouse’ (e.g., stranger in approach) versus the compartment with the newly added stranger mouse.

#### Motor

##### Marbles Buried

The percentage of marbles (n=14) >=75% covered by bedding.

##### Grooming Duration

The total duration (seconds) an animal spent self-grooming.

##### Number of Grooming Epochs

The total number of separate grooming epochs. A grooming epoch was considered if it was >1 second in duration and >1 second separated from the previous epoch. Exploration Velocity: Computed as the average velocity (pixels/frame) of an animals’ centroid during the habituation period of the adult three chamber assay experiments.

##### Chamber transitions

Computed as the total number of times an animals’ centroid crossed from one of the three chambers to another during the habituation, social approach, and social novelty experiments (collectively).

#### Developmental Milestones

##### Self-righting reflex

The first day the reflex was apparent.

##### Eye Opening

The first day both eyes were open.

##### Weaning Weight

Weight at PND 20.

### Phenotype and Neurotype Axes

Phenotype and neurotype axes were defined as the vector normal to the hyperplane that maximally separated VPA and saline animals. To prevent overfitting, hyperplanes were defined by the average beta weights of 5 cross-validated hyperplanes fit on a random 50% folds of the animals. As only a subset of animals was used in imaging experiments, larger folds (60%) of the data were used to estimate the hyperplane. Hyperplanes were fit using MATLAB *fitsvm* function. Behavioral and motif data was scaled (i.e. z-scored across animals) prior to fitting.

An animal’s position along each axis was given by:

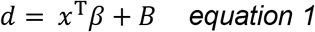

Where *x*^T^ is the transpose of that animal’s 1×12 vector of the behavioral measures or percent explained variance captured by each motif (i.e. motif activity), *β* is a 12 x 1 vector of hyperplane beta weights, and *B* is the shift of the hyperplane relative to the origin (i.e. bias).

### Widefield Imaging Preprocessing

Image stacks were cropped to a 540×540 pixel outline of the cortical window and aligned across sessions using user-drawn fiducials (sagittal sinus and bregma). Bregma was used to align the anatomical reference parcels overlaid in Figures 2. These overlays were created by manually tracing a 2D projection of the Allen Brain Atlas, version CCFv3^34^ (tracing performed with Inkscape Vector Graphics Software). To mitigate potential confounds in our widefield signal due to hemodynamic fluctuations^35–37^, we used both automated and manual masking to conservatively remove non-neural pixels^23^. These masked pixels were ignored during subsequent spatial binning to 68×68 pixels (~136μm^2^/pixel). For each pixel, neural activity was computed as change in fluorescence, e.g. ΔF/F, over time: 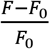. Baseline fluorescence, F0, was computed as the rolling mean of a 9750ms window (130 timepoints) across the entire recording duration. No difference in total variance of neural activity or number of imaged pixels were observed between groups (Extended Data Fig. 7C-D).

Next, we performed pixelwise deconvolution of the ΔF/F signal into an estimate of neural activity using lucy-richardson deconvolution *(lucid* function from^38^; kernel parameters: gamma=0.95, smt=1, p_num=30). This better estimates the underlying neural activity^39^. Previously, we isolated significant activity using filtering and thresholding^23^; deconvolution performs the same function while minimizing data loss. After deconvolution, pixel values were normalized between zero and one to enable CNMF.

Paired-imaging recordings followed the same preprocessing steps.

### Computing Functional Connectivity

Functional connectivity was computed as the fisher r-to-z transformed pairwise correlations between neural activity (ΔF/F) of a grid of regions of interest (ROI) within each cortical hemisphere (Fig. 1F-G). For each ROI, neural activity was taken as the mean deconvolved fluorescence over time of a 9 pixels square centered on that ROI.

### Identifying Spatiotemporal Motifs

We used a custom version of the *seqNMF* algorithm to discover spatiotemporal motifs in widefield imaging data (adapted from *seqNMF* MATLAB toolbox^40^; changed to remove unused features and increase fitting speed). This method employs convolutional non-negative matrix factorization (CNMF) with penalty terms to facilitate discovery of repeating sequences. The general process for motif discovery and clustering across animals is detailed in our previous manuscript^23^. For clarity, we summarize the approach below and detail a few changes in parameter identification.

CNMF approximates the full (pixel by time) image sequence *X_pt_* as a sum of serial convolutions between a spatiotemporal tensor *W* and its temporal weighting matrix *H*:

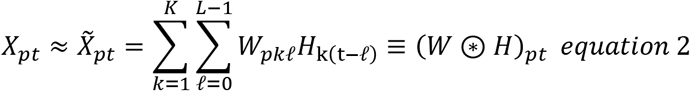

where *K* and *L* are the maximum number of motifs and the maximum length of each motif, respectively and ⊛ indicates the convolutional operator. The sets of motifs in *W* and their temporal weightings in *H* were found with an iterative multiplicative update algorithm.

The value of L was set to 13 frames (975ms). Motifs shorter than this duration were zero padded. This is above the duration of GCaMP6f event kinetics and matches the duration of spontaneous neural events found in previous work^23,41^. The value of K was set to 30 motifs. This was higher than the maximum number of motifs discovered in any fit (Extended Data Fig. 3E).

The optimal spatiotemporal penalty term *λ* of seqNMF was taken as the value that optimized the tradeoff between quality of fit, motif overlap, and number of discovered motifs. This was automatically determined for each fit by sweeping *λ* across 6 orders of magnitude and identifying the cross-over point between quality of fit and motif overlap (Extended Data Fig. 3C). Additional orthogonality and sparsity parameters were held constant for all fits and are reported in Extended Data Table 2.

The multiplicative update algorithm used for CNMF performs poorly on large datasets^42^. Therefore, we fit motifs to two-minute epochs of data from each twelve-minute session. Random initialization leads to variability in identified motifs across fits, so each fit was repeated 10 times. The best fit was taken as the fit with the lowest value of a relative AIC-like measure based on the magnitude of variance in the residuals (which were normally distributed):

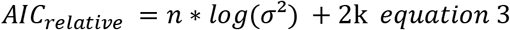

Where n was equal to the number of data points (e.g. *pixels x time*) and *k* was the number of discovered motifs. This measure was used instead of other measures (e.g. explained variance) to avoid selecting fits that captured additional noise.

In total, 105 recordings from 20 animals yielded 4594 single motifs.

To identify motifs across animals, we clustered the single motifs using an unsupervised graph-based approach (Phenograph^43,44^). Phenograph generates a directed graph, where each node (here, an single motif) is connected to its *k* (10) nearest neighbors. Distance between motifs was computed as the peak in their temporal cross correlation. Louvain community detection is then performed on this graph to group nodes into clusters. Prior to clustering, single motifs were smoothed with a 3D gaussian kernel (MATLAB *imgaussfilt3;* σ=[1, 1, 0.1]). Manual inspection revealed 8 of the identified clusters captured imaging artifacts (e.g. hair in imaging field, Extended Data Fig. 3G, inset) and were excluded from subsequent analyses. Collectively these noise clusters included only 182 (4%) of the identified motifs.

The overall (i.e. ‘shared’) motifs fit to all animals were estimated as the mean of the core community of single motifs in each cluster. The core community was defined as the motifs in each cluster that had the most within-cluster nearest neighbors. For a given cluster, the size of this fraction was swept from 1% to 100% of the cluster (5% increments). The optimal fraction was chosen as the value that produced an average motif with the strongest average correlation to all the underlying single motifs in that cluster. In this way, taking the average of the core community prevents averaging-out of the true spatiotemporal dynamics of each cluster. This is necessary because the convolutional nature of seqNMF can lead to motifs in the same cluster having slight temporal jitter and warping.

Prior to averaging, the motifs in the core community were aligned to a ‘template’ motif. The template motif shared the most zero-lag peak temporal cross-correlations with all other motifs within that cluster. If there were multiple templates with equivalent cross correlations, then one was chosen at random. All motifs were zero-padded to a length of *3L* (39 timepoints) to allow for some temporal jitter and aligned by maximizing their cross-correlation lag to the template motif from their cluster. The average overall motif for the cluster was then estimated by averaging the activity of the temporally aligned core community. Finally, we aligned the motifs to one another by shifting the center of mass of activity to the middle timepoint. Timepoints with no variance across all overall motifs were removed. This removed frames without any neural activity in any motif and resulted in motifs with a duration of 27 timepoints (~2s).

The resulting 16 shared motifs identified across all animals were then refit to the each 2-minute epoch of data using the same seqNMF algorithm. Here, *W* (e.g. the tensor of motifs) was fixed to prevent changes in the motifs while allowing their temporal weights (*H*) to be optimized. Prior to refitting, the original data were spatially smoothed with a 2D gaussian filter. The sigma of this filter was autofit for each epoch to best match the average spatial variance of the motifs (bounded between σ=[0, 0] and [1.5, 1.5]; median autofit value=[0.75, 0.75]). This same refitting procedure was used to refit the 16 motifs to the paired-imaging data.

While similar to our original approach for estimating motifs^23^, the approach outlined here introduces improvements in preprocessing (e.g. deconvolution), new automated approaches for parameter selection, and the addition of 11 new (VPA) animals to our analysis. Despite these changes, the number and structure of identified motifs was highly similar to our previous work (e.g. ~16 motifs shared across animals explaining ~75% of the variance in neural activity; see Table S1 for comparison of motifs between studies). The changes in preprocessing did result in fewer motifs discovered per 2-minute epoch (Extended Data Fig. 3E), likely due to less spatial noise in deconvolved data (verse the previously thresholded data) and the AIC-like criterion selecting more parsimonious individual fits.

### Linear Classification to Compare Motifs

Linear classifiers with leave-one-out cross-validation were used to test whether the spatiotemporal structure of motifs differed between groups. For each of the 16 shared motifs, the single motifs from one animal that contributed to that shared motif were withheld, and a classifier was trained to differentiate between saline and VPA animals using the contributing motifs from all other animals (balanced between groups). The classifier was then tested to determine the group identity of the motifs from the withheld animal. This was repeated for all animals for each of the 16 overall motifs. These classifiers were unable to accurately assign group labels above chance (see Main text for statistics). Multiple classifier types were tested, including linear discriminant, support vector machine (svm) with standard linear kernels, svm with radial basis function kernels, svm with Bayesian-optimized hyperparameters, and svm with ANOVA-based feature selection. All failed to accurately distinguish motifs between groups.

To confirm classifier efficacy, a similar classification strategy was used to compare between motifs. For all 120 n-choose-two combinations of the 16 shared motifs, the contributing motifs from one animal were withheld and a classifier trained to identify the motif label of the motifs from the remaining animals. Classifiers were then tested to determine the label of the motifs in the withheld animal, which they did with high accuracy (see Main text for statistics).

### Percent Explained Variance Calculations

Reconstructions of the original data were computed by convolving motifs with their temporal weightings (equation 2). Percent variance in neural activity captured by these reconstructions were computed as

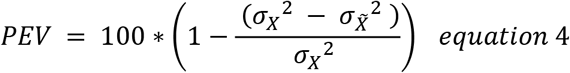

Where *σ_X_*^2^ and 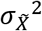 denote the spatiotemporal variance of the original data and reconstructed data respectively. Relative PEVs, referred to as the ‘activity’ of single motifs, were calculated by dividing the PEV of each motif by the total PEV across all motifs for that epoch. These values were averaged across epochs and recordings for individual mice to yield a single value per motif per mouse.

Motif activity in paired context (Fig. 3) was calculated the same way and the change in motif activity from alone to paired contexts was measured as the sum squared error between motif activity in each context. To determine if change in motif activity was greater than chance, a null distribution was estimated for each animal by randomly permuting the variance captured by motifs between the two contexts.

## Extended Data Figures and Tables

**Extended Data Figure 1:**
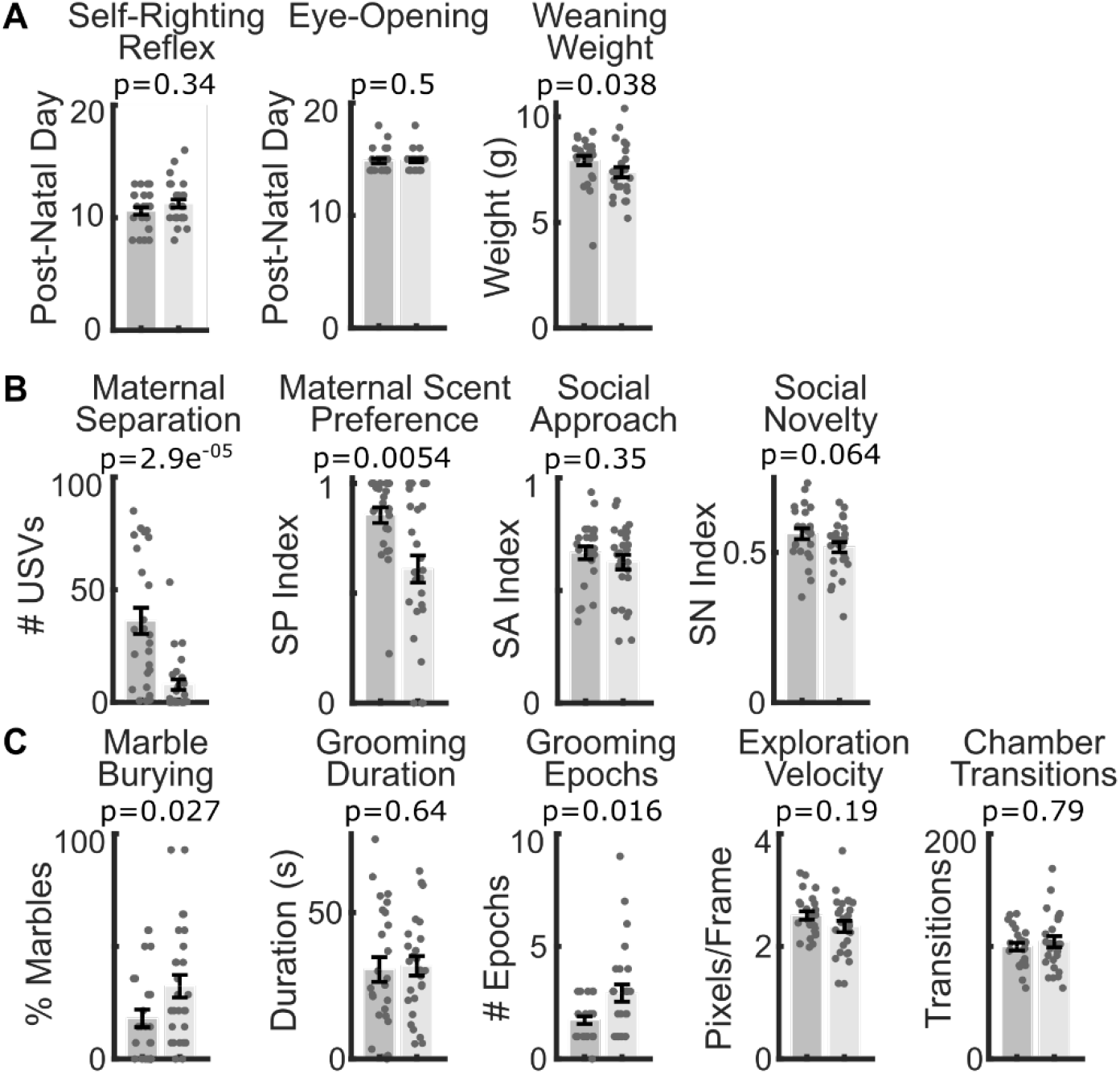
Results of individual behavioral tests. **(a-c)** Results comparing behavior of mice exposed to valproic acid (VPA) *in utero* to that of saline-exposed animals. Tests are grouped by (a) developmental, (b) sensory-memory, and (c) motor behavioral domains. Full distribution and mean +/- SEM shown, p-value estimated with Mann-Whitney U-Test.

**Extended Data Figure 2:**
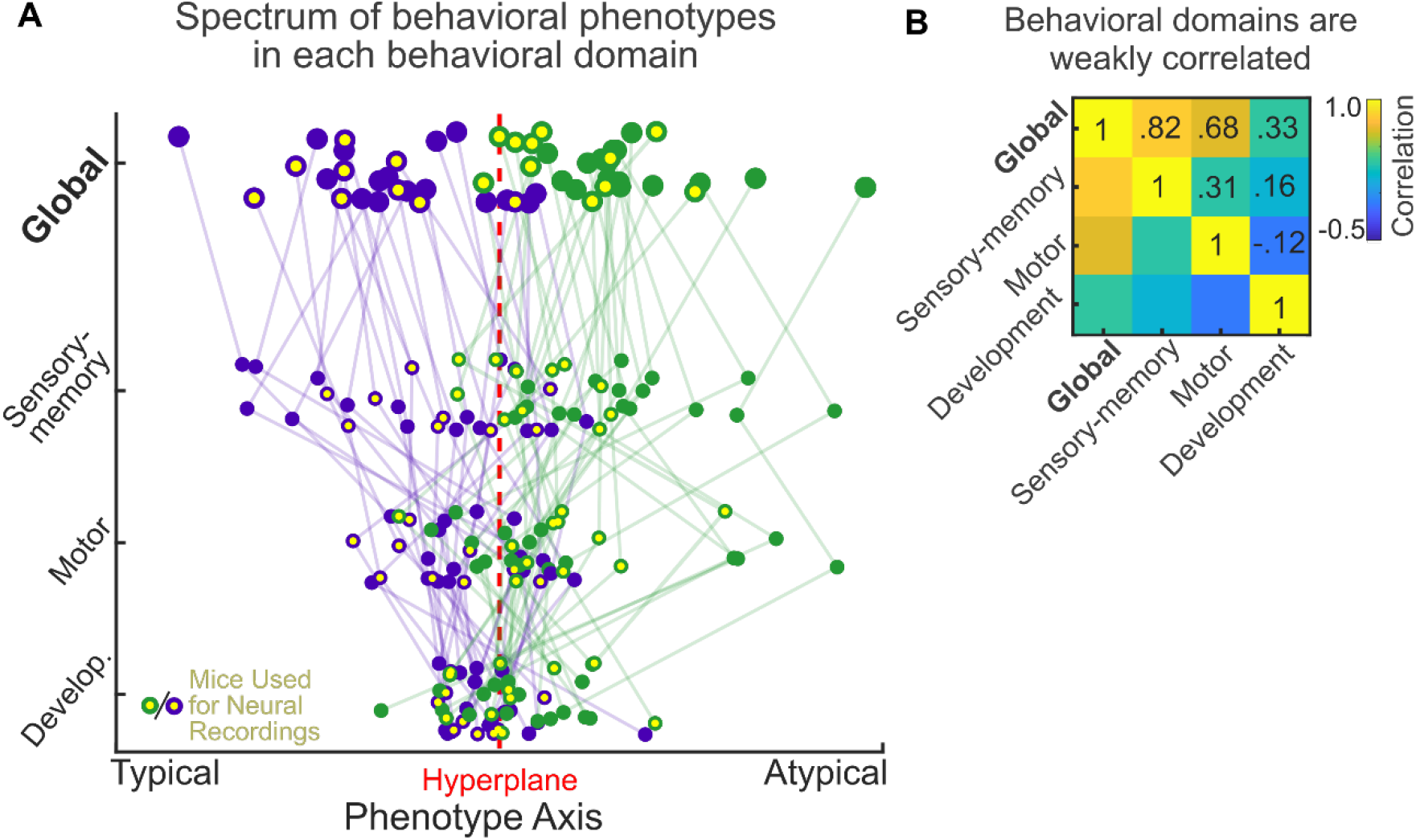
Phenotype axes in each behavioral domain.

**Extended Data Figure 3:**
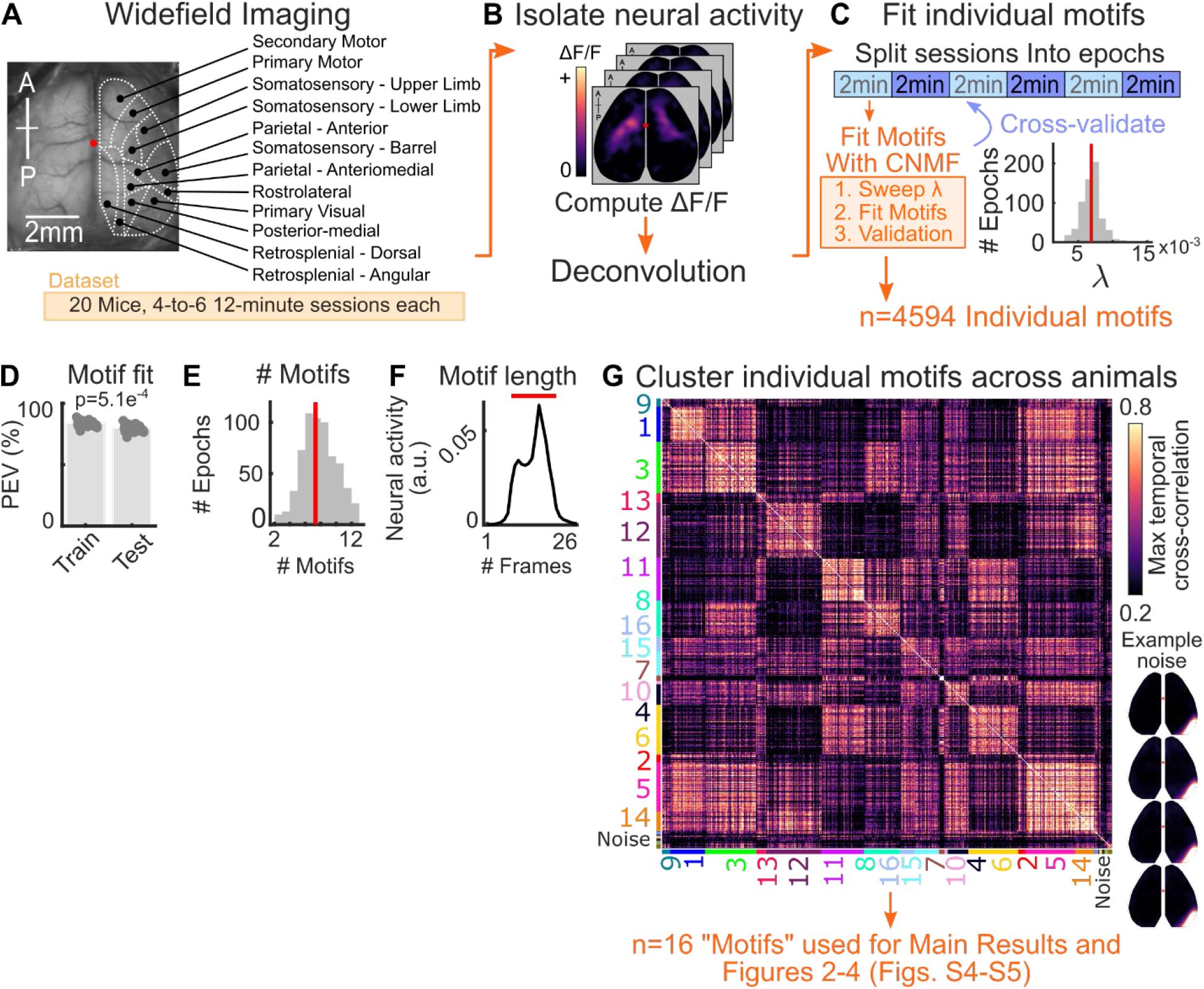
Identifying spatiotemporal motifs in cortical neural activity. Pixel value reflects the relative neural activity within the motif over time. Units are arbitrary as the true pixel value is the result of motifs convolved with their temporal weightings (see Methods). Pixel activity was averaged within each motif and then averaged across all n=4594 motifs. Prior to averaging, motifs were zero-padded and aligned by their maximum autocorrelation. Importantly, the timecourse of motif activity rose and fell within the 13-frame maximum duration of each motif (red line shows length of 13 frames), suggesting that this parameter did not limit the identified motifs (see Methods). **(g)** Pairwise peak temporal cross-correlation between all 4594 identified motifs. Motifs are grouped by their membership in motif clusters. Cluster identity is indicated with color code along axes. Group numbering (and thus the sorting of the correlation matrix) is determined by beta values of each motif in Figure 2D. Both VPA and saline animals contributed to each cluster. Manual inspection removed 8 clusters corresponding to non-neural activity (i.e. lateral hair intrusion; labeled ‘noise’; see inset for example). All p-values estimated with Mann-Whitney U-Test.

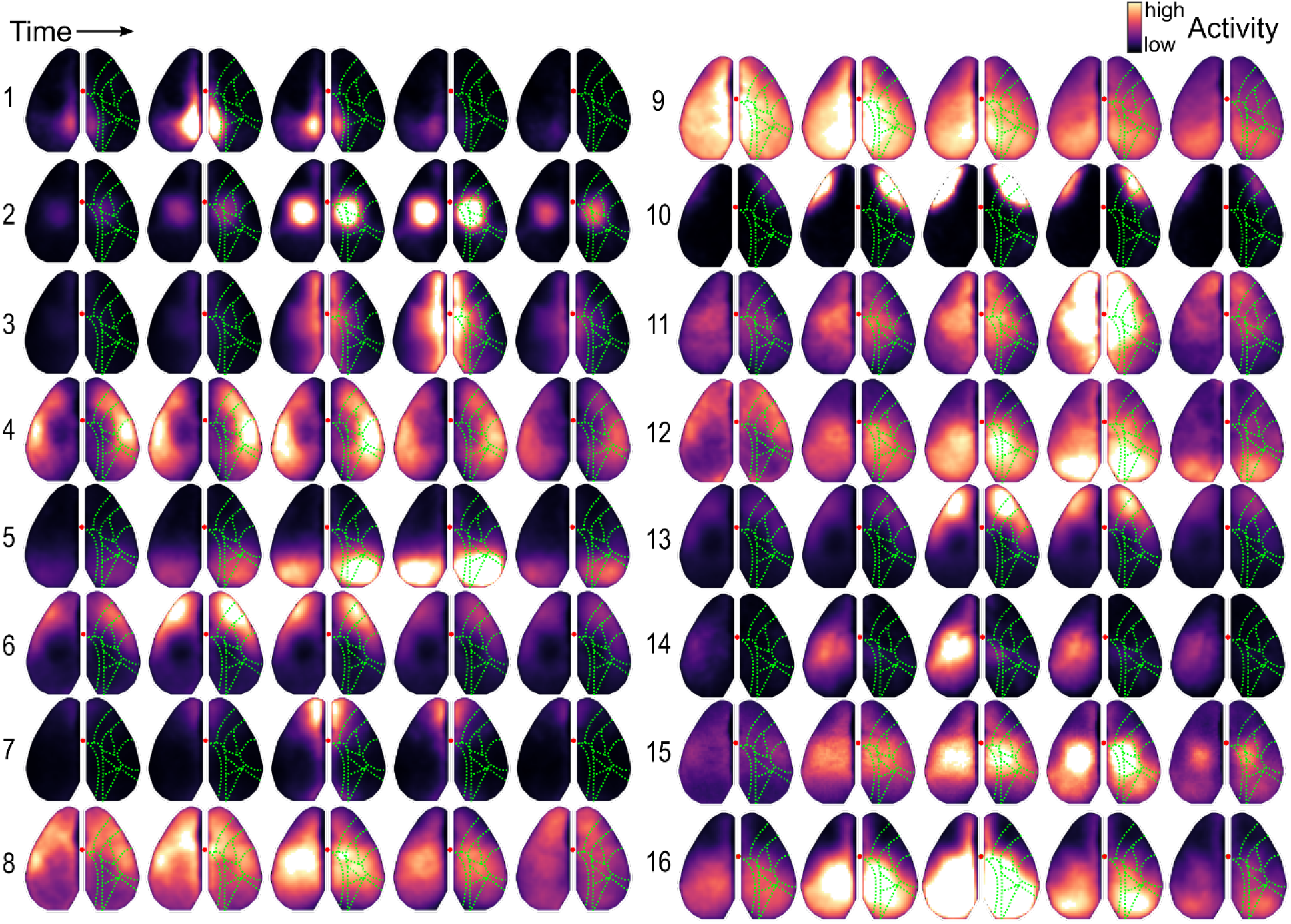
Display follows Figure 2C. Saturation is scaled per motif. Motifs are ordered by beta weights in Figure 2D. Written descriptions are provided in Table S1.

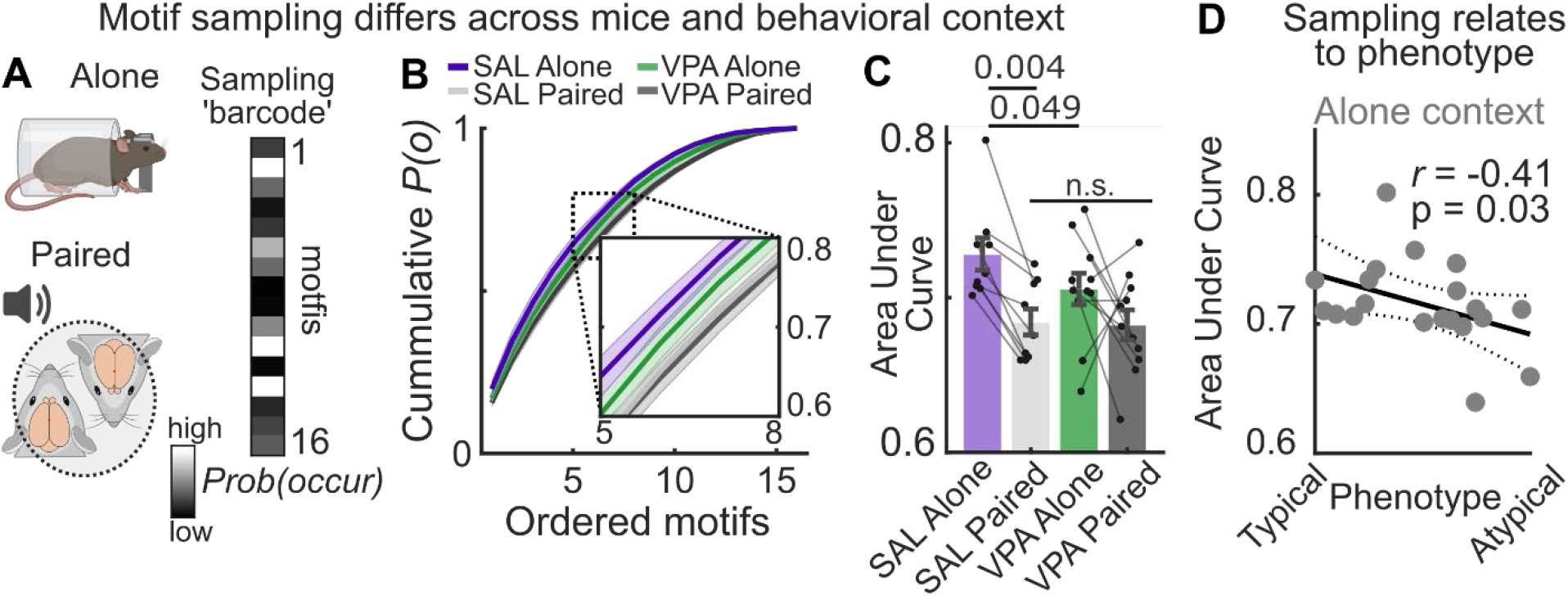
**(a)** How often each motif occurred was measured in two behavioral contexts; when mice were alone, at rest, or paired with another animal in a stimulus-rich environment (see Methods). Motif activity in either environment was used to define ‘barcodes’ in motif sampling. **(b)** Cumulative probability of motifs occurring as a function of motifs ordered by descending probability of occurrence (per animal). If motifs are sampled to match a given environment, then the probability of certain motifs occurring will be greater than others (i.e., less uniform barcode), resulting in a steep curve in the cumulative probability distribution across motifs and a large area under this curve. In contrast, a flatter curve and lower AUC would indicate that motifs occur more uniformly. Light gray “SAL Paired” line is behind dark gray “VPA Paired” line. **(c)** Area under curve of cumulative probability distribution across groups and contexts. Consistent with the hypothesis that motifs are sampled to match the environment, motifs occurred significantly less uniformly in all saline animals when alone compared to when in the novel stimulus-rich paired context (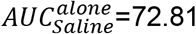, CI: 71.46-75.85 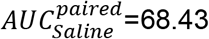, CI: 66.95-70.02; p=0.004, Wilcoxon signed-rank test). However, motif sampling differed between groups and was related to behavioral phenotype. In the alone context, motifs occurred more uniformly in VPA mice than saline mice (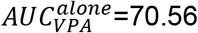, CI: 68.43-72.16; p=0.049, permutation test). **(d)** Area under curve as a function of animal phenotype in the alone context. Across mice, uniformity in motif sampling barcodes was significantly negatively correlated with behavioral phenotype. In other words, more atypical animals more broadly sampled different motifs when alone, at rest. For all plots, data points show individual mice, error bars/distributions reflect mean +/- SEM. Solid and dotted lines in panel D show fit and 95% confidence bounds, respectively.

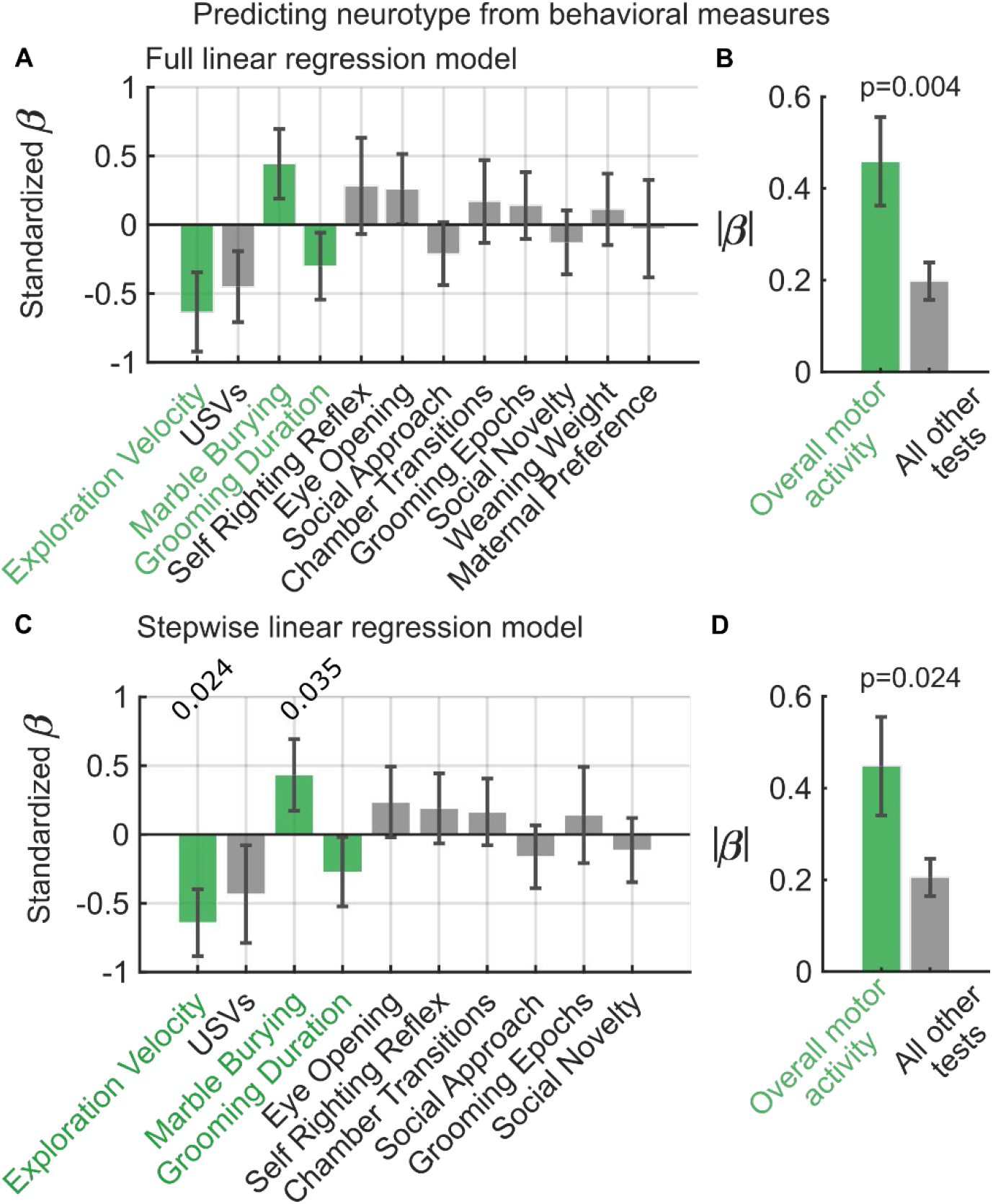
**(a)** Multiple linear regression was performed to test the association between the results of 12 behavioral measures in each animal (i.e., Extended Data Fig. 1a-c) and animals’ neurotype (i.e., cortical motif activity, Fig. 2e; see Methods). Bars show estimated coefficients (β; y-axis) for each behavioral measure (x-axis). Error bars show standard error. Behavioral measures are ordered by decreasing absolute value of β, revealing that measures that captured overall motor activity were most strongly associated with animals’ neurotype (green; these measures were exploration velocity, marble-burying, and total self-grooming time). **(b)** Measures of gross motor activity were significantly more strongly associated with neurotype than other behavioral tests (p=0.004, permutation test). Plot shows the average absolute value of βs for gross motor activity measures (green; left) and the remaining measures (gray; right). Bar and error bars show mean and standard error, respectively. **(c-d)** Stepwise linear modeling confirmed that potential overparameterization in the full model (panels a and b) did not impact findings. Stepwise regression showed that multiple behavioral tests, particularly those associated with gross motor phenotype, were significantly predictive of neurotype. Presentation follows panels a and b. Full model fit: R^2^= 0.81 p=0.12. Stepwise model fit: R^2^= 0.8 p=0.032.

**Extended Data Figure 7:**
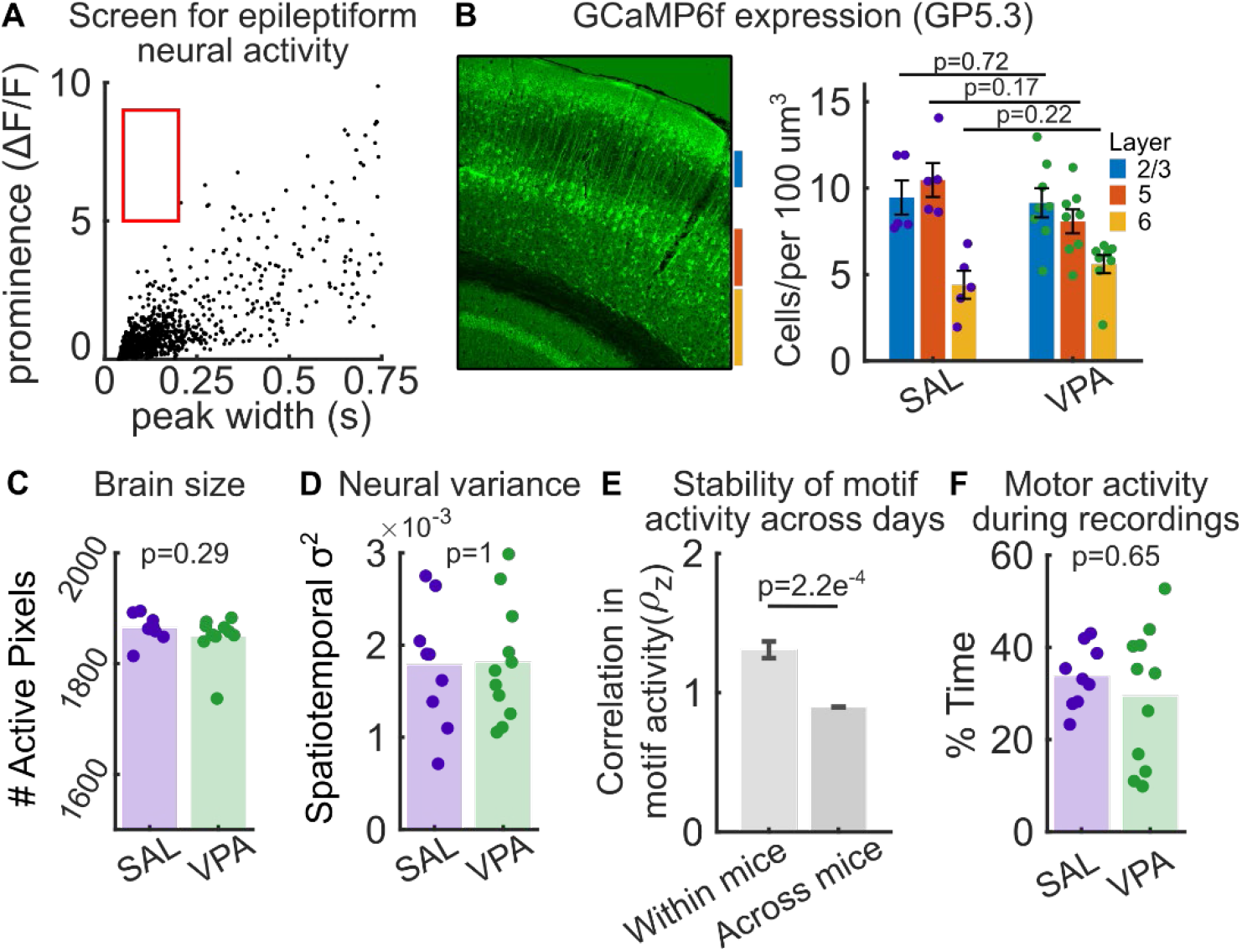
Additional control experiments. **(a)** No animal exhibited evidence of the epileptiform events reported in other GCaMP6f lines. Plot shows post-hoc screening for epileptiform events as in^26,28^. Epileptiform cortical activity in widefield imaging data is characterized by a cluster of high amplitude, short duration events (red square). No such clusters were observed in any animal. **(b)** Left: Example confocal microscopy of GCaMP6f (green) expressing cells in a coronal section of the primary visual cortex. Right: Manual cell counting confirmed no differences in GCaMP6f expression between VPA and saline mice across different cortical layers. Cells were counted using *Fiji* by an experimenter blinded to animal groups. Full distribution and mean +/- SEM shown **(c)** Confirmation of similar sized mesoscale imaging field of view between groups. Field of view was quantified as the number of pixels with non-zero variance after masking non neural pixels. Full distribution and mean shown. **(d)** Confirmation of similar total amounts of neural activity between groups. Neural activity was quantified as the spatiotemporal variance (variance of the vectorized image series) of activity over time. Variance was calculated on preprocessed and deconvolved data and averaged across sessions for each animal. **(e)** Confirmation that variance in motif activity was more stable within animals than between animals across recording days. Stability was quantified as the average correlation between the vector of motif percent explained variances (i.e., the barcode of motif activity) across days (within-mice) or across animals (across-mice). Correlations show fisher r-to-z transformed Pearson correlation coefficients. **(f)** Confirmation that motor activity during recordings was similar between groups. Gross locomotor activity was measured by a piezo motion sensor placed on an animal’s back. ‘Active’ periods were quantified by thresholding the voltage readout of the piezo. Plots show average percent of total recording time spent in active periods per mouse. In addition, the amount of motor activity during a recording was not correlated with animals’ neurotype (r=0.08, p=0.65). Full distribution and mean shown. All p-values estimated with Mann-Whitney U-Test or Wilcoxon Signed-Rank Test (panel e).

**Extended Data Figure 8:**
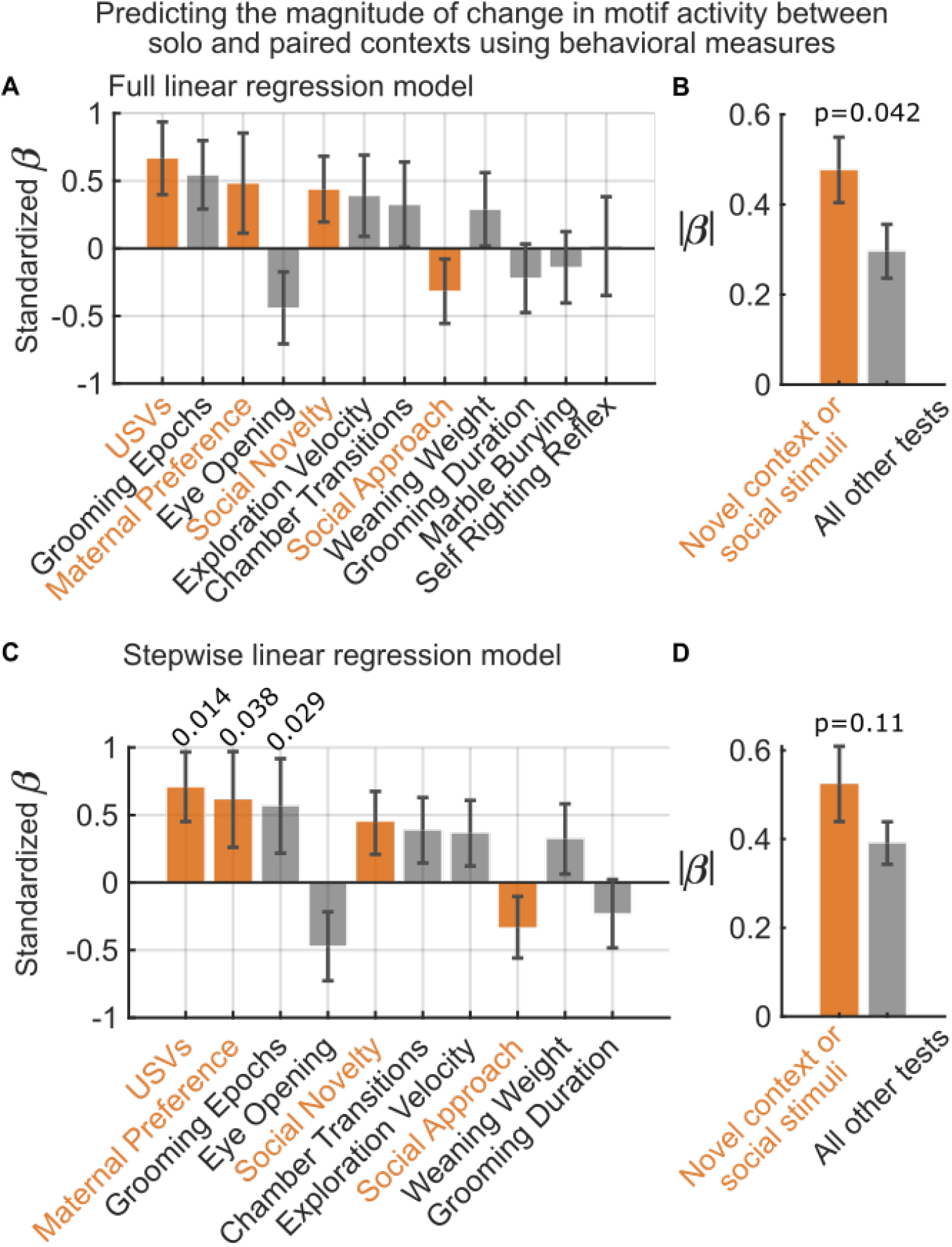
Evaluating the relationship between behavioral measures and the magnitude of change in motif sampling between alone and paired contexts. **(a)** Multiple linear regression was performed to test the association between the results of 12 behavioral measures in each animal (i.e., Extended Data Fig. 1a-c) and the magnitude of change in animals’ motif activity between the alone and paired contexts (i.e., the SSE in motif ‘barcodes’ between contexts from Fig. 3a-b). Bars show estimated coefficients (β; y-axis) for each behavioral measure (x-axis). Error bars show standard error. Behavioral measures are ordered by decreasing absolute value of β. Interestingly, many of the behavioral measures from tests evaluating the sensory-memory domain – which includes tests of sociability and/or novel contexts – were amongst the most strongly associated with animals’ motif activity (orange). These measures were pup ultrasonic vocalizations (USVs) in response to maternal separation, juvenile preference for maternal scent, and three-chamber social novelty and social approach. **(b)** Collectively, these measures were significantly more strongly associated with neurotype than other behavioral tests (p=0.042, permutation test). Bar plot shows the average absolute value of βs for these measures (orange; left) and the remaining behavioral tests (gray; right). Bar and error bars show mean and standard error, respectively. **(c-d)** Stepwise linear modeling confirmed that potential overparameterization in the full model (panels a and b) had minimal impact on results. Stepwise regression showed that multiple sensory-memory behavioral measures were significantly predictive of changes in motif sampling. More generally, a similar trend was observed in which these tests were amongst the most strongly associated with animals’ motif sampling and trended towards stronger βs relative to other behavioral measures. Presentation follows panels a and b. Full model statistics: R^2^= 0.79, p=0.15. Stepwise model statistics: R^2^=0.78, p=0.046

**Extended Data Figure 9:**
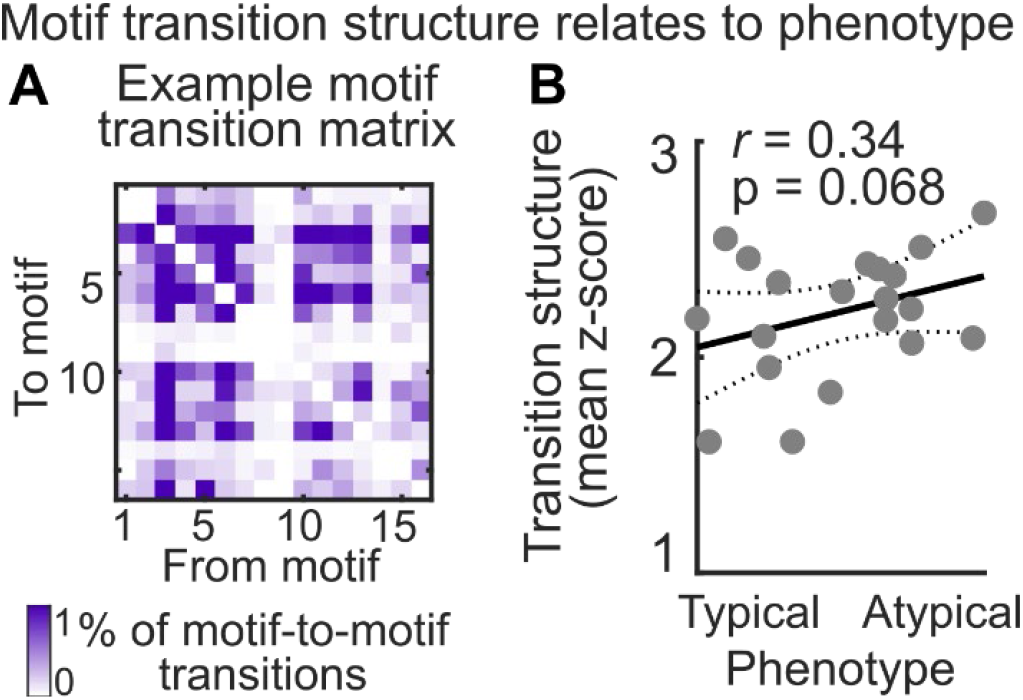
Investigating transitions between motifs over time. **(a)** Transition matrix from an example saline animal showing the probability of each possible transition between motifs over time (excludes self-transitions). **(b)** Magnitude of structure in the motif transition matrix compared to behavioral phenotype across animals in the alone context. For each animal, the probability of a given motif transition was z-scored relative to a null probability distribution for that transition estimated by discrete Markov chain modelling. The absolute z-scored transition probabilities were averaged per animal, giving a measure of the magnitude of transition structure relative to chance, i.e., higher values correspond to more structured and less stochastic transitions between motifs. Across animals, transition structure trended towards positively correlating with phenotype. In other words, more atypical animals showed more stereotyped transitions between motif over time.

**Extended Data Table 1.** Detailed description of each motif.

| # | Ordered List of Areas Activated | General Description | *Most Similar Motif # in |
| --- | --- | --- | --- |
| 1 | RSP | Burst in activity in retrosplenial cortex. | 12 |
| 2 | PC | Localized burst in parietal regions. | 3 |
| 3 | [medial M2, RSP] | Burst of activity in medial regions. | 7 |
| 4 | [BC, RL, SS, anterolateral M1/M2] → [BC, RL, amPC, anterior VC] | Posteromedial wave of activity from barrel cortex and anterolateral motor cortices to visual areas. | ~9 |
| 5 | [V1, PM, posterior RSP] | Localized burst in visual areas. | 10 |
| 6 | [anterolateral M1/M2, SS, BC] | Burst of activity in anterolateral somatosensory, motor, and barrel cortex *** | 6 |
| 7 | M2 | Burst in activity of secondary motor cortex. | 13 |
| 8 | [anterolateral M1/M2, SS, BC] → PC → M2 | Anterior-to-posterior wave from motor and somatosensory regions to parietal cortex, followed by a posterior-to-anterior wave from parietal to secondary motor cortex. | ** |
| 9 | [medial M2, PC, anterior RSP] → [RSP → [RSP, V1] | Anterior-to-posterior midline wave of activity across the cortex. | ~1 |
| 10 | [lateral M1/M2, SS, BC] | Burst of activity in lateral motor and somatosensory areas. | 14 |
| 11 | PC → anteromedial M1/M2 → [RSP, PC] | Activity in parietal cortex followed by an anterior-to-posterior wave of activity from medial motor cortex to retrosplenial areas. | 8 |
| 12 | [anterolateral M1/M2, SS, BC] → [medial M1/M2/SS/BC] → PC → [RSP, PC, V1] → V1 | Anterior-to-posterior wave of activity across the cortex. | 11 |
| 13 | [anterolateral M1/M2, SS, BC] → [anteromedial M1/M2, SS] | Wave of activity from anterolateral somatosensory, motor, and barrel cortex to anteromedial somatosensory and motor areas.*** | 6 |
| 14 | [left PC] | Largely unilateral burst of activity in left parietal cortex | ** |
| 15 | [RL,BC] → PC → [PC, VC] | Rostrolateral and barrel cortex activity followed by activity in parietal and visual regions. | 5 |
| 16 | [RSP, PC] → [RSP, PC, VC] → [VC] | Posterolateral wave of activity from retrosplenial and visual areas. | 4 |

[ ] denotes co-activated areas. → denotes sequential activation.
SM = secondary motor, PM = primary motor, SS = somatosensory, BC = Barrel Cortex, PC = parietal cortex (all), amPC = anterior-medial parietal cortex, aPC = anterior parietal cortex, RL = rostrolateral cortex, VC = primary visual cortex, RSP = retrosplenial cortex (all).
\* Comparisons are qualitative. Many motifs observed in both studies were highly similar. Some differences in single motifs were observed; likely due to clustering across ~2x number of motifs (and mice) and differences in preprocessing of the data (see Methods).
\*\* Motif not observed in<sup>23</sup>.
\*\*\* Motifs 6 and 13 are qualitatively very similar. However, they exhibit different activity in the data (Fig. 2D), suggesting that they reflect different patterns. Indeed, motif 6 presents more as a wave, while motif 13 presents as a burst.

**Extended Data Table 2.**
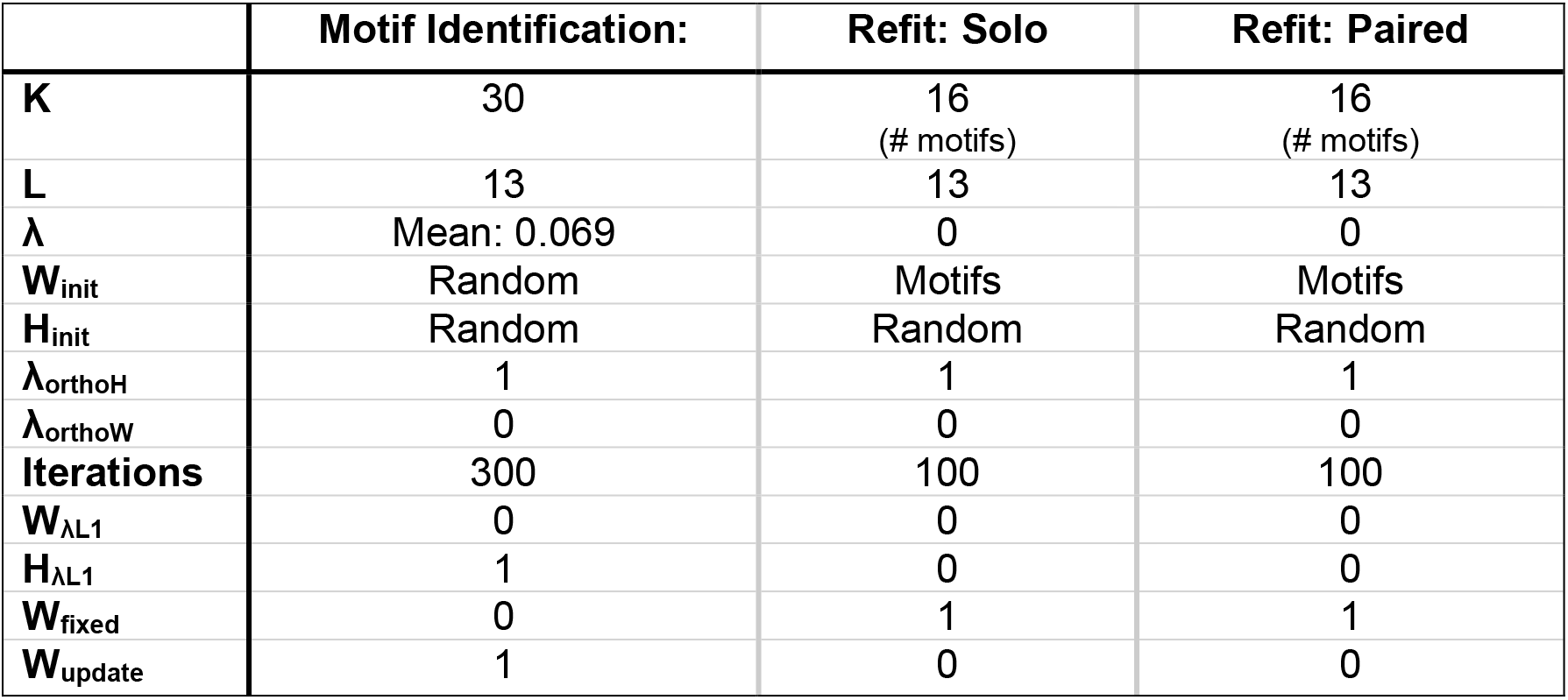
CNMF parameters. See Methods for details of parameter choices. Parameters are labeled as in original seqNMF toolbox^40^. Analysis code used an adapted version of this toolbox. In this adapted version, λ is 100 times larger (e.g. seqNMF λ would equal 0.00069).

